# From thoughtless awareness to effortful cognition: alpha - theta cross-frequency dynamics in experienced meditators during meditation, rest and arithmetic

**DOI:** 10.1101/2020.01.14.905935

**Authors:** Julio Rodriguez-Larios, Pascal Faber, Peter Achermann, Shisei Tei, Kaat Alaerts

**Affiliations:** University of Leuven, KU Leuven, Belgium, Department of Rehabilitation Sciences, Research Group for Neurorehabilitation; The KEY Institute for Brain-Mind Research, Department of Psychiatry, Psychotherapy and Psychosomatics, University Hospital of Psychiatry, Zurich, Switzerland; Institute of Pharmacology and Toxicology, University of Zurich, Winterthurerstrasse 190, 8057 Zurich; Neuroscience Center Zurich, University of Zurich and ETH Zurich, Zurich, Switzerland; Zurich Center for Interdisciplinary Sleep Research, University of Zurich, Zurich, Switzerland; Department of Psychiatry, Kyoto University Graduate School of Medicine, Sakyo-ku, Kyoto 606-8507, Japan

## Abstract

Neural activity is known to oscillate within discrete frequency bands and the synchronization between these rhythms is hypothesized to underlie information integration in the brain. Since strict synchronization is only possible for harmonic frequencies, a recent theory proposes that the interaction between different brain rhythms is facilitated by transient harmonic frequency arrangements. In this line, it has been recently shown that the transient occurrence of 2:1 harmonic cross-frequency relationships between alpha and theta rhythms (i.e. f_alpha_≈12 Hz; f_theta_≈6 Hz) is enhanced during effortful cognition. In this study, we tested whether achieving a state of ‘mental emptiness’ during meditation is accompanied by a relative decrease in the occurrence of 2:1 harmonic cross-frequency relationships between alpha and theta rhythms. Continuous EEG recordings (19 electrodes) were obtained from 43 highly experienced meditators during meditation practice, rest and an arithmetic task. We show that the occurrence of transient alpha:theta 2:1 harmonic relationships increased linearly from a meditative to an active cognitive processing state (i.e. meditation< rest< arithmetic task). It is argued that transient EEG cross-frequency arrangements that prevent alpha:theta cross-frequency coupling could facilitate the experience of ‘mental emptiness’ by avoiding the interaction between the memory and executive components of cognition.

## INTRODUCTION

Despite the heterogeneity of meditation traditions and styles, there seem to be some common denominators in their objectives, techniques and phenomenology. In this way, an important part of meditation practices are aimed at achieving an equanimous mind state of thoughtless awareness^1–3^, by instructing meditators to focus on an object (e.g. breathing, bodily sensations) and to bring the attention back to the present moment (non-judgementally) whenever mind-wandering episodes occur^4^. These kind of present-centered strategies have been shown to decrease mind-wandering frequency and engagement thereby leading to the subjective experience of ‘non-involvement’, ‘detachment’ or ‘ego dissolution’^5–9^. Because, from a cognitive perspective, self-generated thought requires retention and manipulation of information^10,11^, it can be anticipated that meditation practices that are aimed at achieving a state of ‘mental emptiness’ (by reducing mind wandering frequency and engagement) involve a diminution of the interaction between the memory (retrieval and retention of information) and executive (manipulation of information) components of cognition.

Brain oscillations at alpha (8 – 14 Hz) and theta (4 – 8 Hz) frequency bands have been shown play to key role in a wide variety of cognitive tasks involving memory and executive control^12^. While alpha oscillations have been associated with the storage and retrieval of information; theta oscillations have been associated with the manipulation of information^13–17^. Accordingly, changes in alpha-theta cross-frequency interactions (depending on cognitive task demands) have been suggested to reflect the degree of integration between the executive (i.e. theta) and memory (i.e. alpha) components of cognition^18–23^.

Effortful cognitive tasks have been shown to induce an acceleration of alpha peak frequency^24,25^ and the emergence of a theta peak around its harmonic (i.e. theta peak ~ alpha peak / 2)^21,26^. Since brain rhythms are thought to transmit information through their synchronization^27–29^ and strict synchrony is only possible between harmonic frequencies^30,31^, it has been recently proposed that communication between the neural networks underlying alpha and theta rhythms is facilitated by the formation of a harmonic frequency arrangement (i.e. theta frequency = alpha frequency /2)^32,33^. In support of this notion, a previous study recently showed that the incidence of ‘2:1’ ‘harmonic’ relationships between the peak frequencies of alpha and theta rhythms (e.g. alpha= 10 Hz; theta= 5 Hz) increases during a cognitively demanding arithmetic task (compared to rest) and correlates with improved arithmetic task performance^34^. Furthermore, and in line with the idea that a ‘2:1’ harmonic relationship facilitates synchrony between two neural oscillators^30,32^, it was also shown that the increased incidence of harmonic alpha-theta cross-frequency arrangements is tightly associated with increased cross-frequency phase synchronisation^34^.

A substantial number of EEG studies have investigated changes in alpha and theta oscillations during meditative states. In this regard, a recent review concluded that, although not uniformly reported, the most consistent finding in the literature is an increase in alpha and theta band power during meditation compared to rest^35,35,36^. However, as highlighted by these authors, the reviewed studies differed significantly with respect to the adopted EEG measurements, the criteria to classify participants as ‘experienced meditators’ as well as the type of meditation practice^35^. In addition to band power modulations, peak frequency changes in the theta-alpha range have also been studied in relation to meditation practices. Here, studies seem to converge on the finding that individual alpha frequency (typically defined as the center of gravity between 7 and 14 Hz; see Klimesch, Schimke, & Pfurtscheller, 1993) decelerates during meditation practice compared to rest^38–41^. Importantly, despite the extensive literature pointing to the importance of alpha-theta cross-frequency dynamics for cognitive processing^20–23,34^, their potential modulation during meditative states remains unexplored.

In this study, we aimed to assess whether the neural correlates of meditative states can be characterized in opposition to active cognitive processing from a cross-frequency dynamics perspective. Specifically, we hypothesized that the incidence of an EEG cross-frequency arrangement that facilitates alpha:theta cross-frequency synchronization would increase linearly from a meditative to an active cognitive processing state (i.e. meditation < rest < cognitive task). In order to test this hypothesis, we estimated the transient occurrence of harmonic relationships between the instantaneous peak frequencies of alpha and theta rhythms (alpha peak frequency / theta peak frequency = 2.0) in a sample of 43 highly experienced meditators that (i) were engaged into their respective meditation practice, (ii) sat in a (non-meditative) resting state, and (iii) performed a cognitively demanding arithmetic task.

## METHODS

### Participants

Data analyses were performed using an EEG dataset of 43 highly experienced meditators from three different meditation traditions (i.e. 15 QiGong, 14 Sahaja Yoga, 14 Ananda Marga Yoga) that were adopted from a larger EEG dataset^42^. The study was approved by the Ethics Committee of the Tokyo University Medical School (#1364) and it was performed in accordance with the relevant guidelines and regulations. The participants were fully informed about the goal and methods of the study and gave their written informed consent. Table 1 depicts the mean age, gender, years of meditation experience and a brief description of the respective meditation objectives for each of the included meditation traditions (for more details see Lehmann et al., 2012).

**Table 1.**
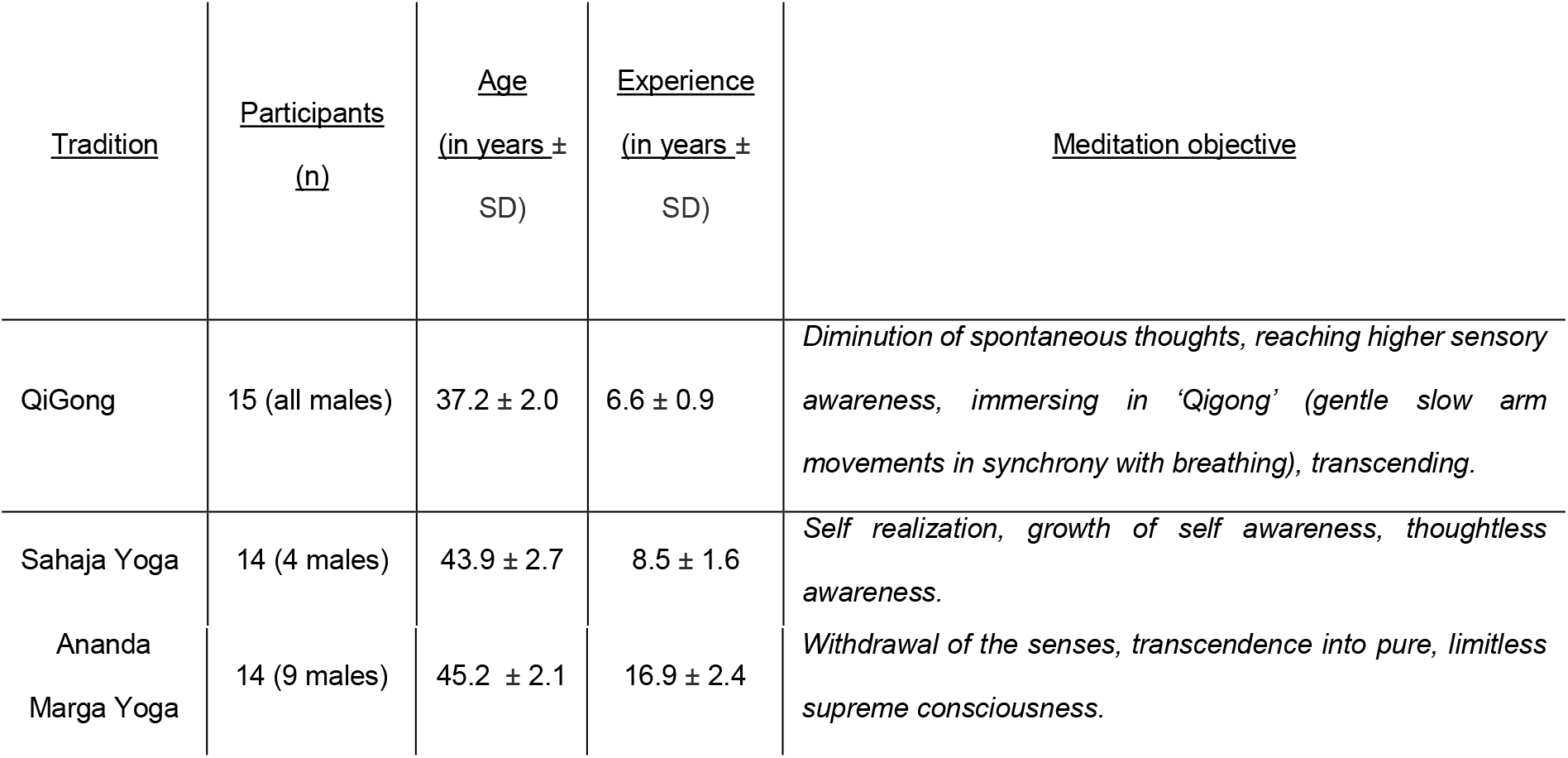
Demographics of the participants.

### Experimental design

The EEG recording session encompassed the following 6 conditions: initial rest, breath counting, meditation, intermediate rest, mental arithmetic task and post arithmetic rest. In the current study, only the initial rest, meditation and mental arithmetic conditions were used for analyses. In the mental arithmetic task, participants were asked to subtract 7 iteratively from 1000 and to re-start from 1000 if they lost track or reached zero for a total duration of 5 minutes. The rest condition consisted of 4 cycles of 20 s ‘eyes open’ alternated with 40 s ‘eyes closed’. Only the ‘eyes closed’ cycles were used for analyses. During the meditation condition, participants were asked to engage into their respective meditation practice for a total duration of 20 min. Note that, only the (approximately) middle 3 minutes of the meditation session (from the pool of available artefact-free epochs) were used for analyses in order to make the length of the different conditions comparable.

### EEG recordings and pre-processing

The EEG recordings were performed in the General Veterans Hospital in Taipei, from September to December 2006. EEG recordings and pre-processing steps were performed as described in Lehmann *et al*.^42^. In short, the EEG was recorded with combined ears as a reference (standard 19-EEG channels with international 10/20 system). Raw EEG signals were band pass filtered from 1 to 70 Hz and digitized at 256 samples/s. Independent component analysis was used to correct for eye movements. Then, data was epoched (2 seconds) in order to manually reject artefactual epochs by visual inspection. Up to 3 bad channels were replaced by the average of the direct neighbouring channels. The mean number of channels that were interpolated was 0.73. For all of the following analyses, the available artefact-free epochs were concatenated.

### Instantaneous frequency and transient numerical ratio calculation

Instantaneous peak frequencies in the alpha (8 - 14 Hz) and theta band (4 – 8 Hz) were computed in MATLAB (version r2017b) using the method and code developed by Cohen^43^ which estimates the instantaneous peak frequencies in a frequency band based on the temporal derivative of the phase angle time series.

First, the EEG signal was filtered for alpha (8 - 14 Hz) and theta (4 - 8Hz) bands with a plateau-shaped zero phase digital filter with transition zones of 15 %^43^. Next, the Hilbert transform was applied to extract the phase angle time series based on the analytical signal. The instantaneous frequency was determined by multiplying the first temporal derivative of the phase angle by the sampling rate and dividing it by 2π. A median filter was applied to the instantaneous frequency time series using the parameters suggested by Cohen^43^ in order to attenuate non-physiological frequency jumps. Finally, the numerical ratio between the instantaneous alpha and theta frequencies was computed per time point and averaged to the first decimal place for the calculation of the proportion (%) of time points showing a harmonic 2:1 relationship (i.e. alpha frequency / theta frequency = 2.0) (see Figure 1A). The proportion (%) of time points showing a harmonic 2:1 relationship (hence forward termed ‘harmonic locking’) was determined separately for each condition (meditation, rest, arithmetic), subject and electrode.

**Figure 1.**
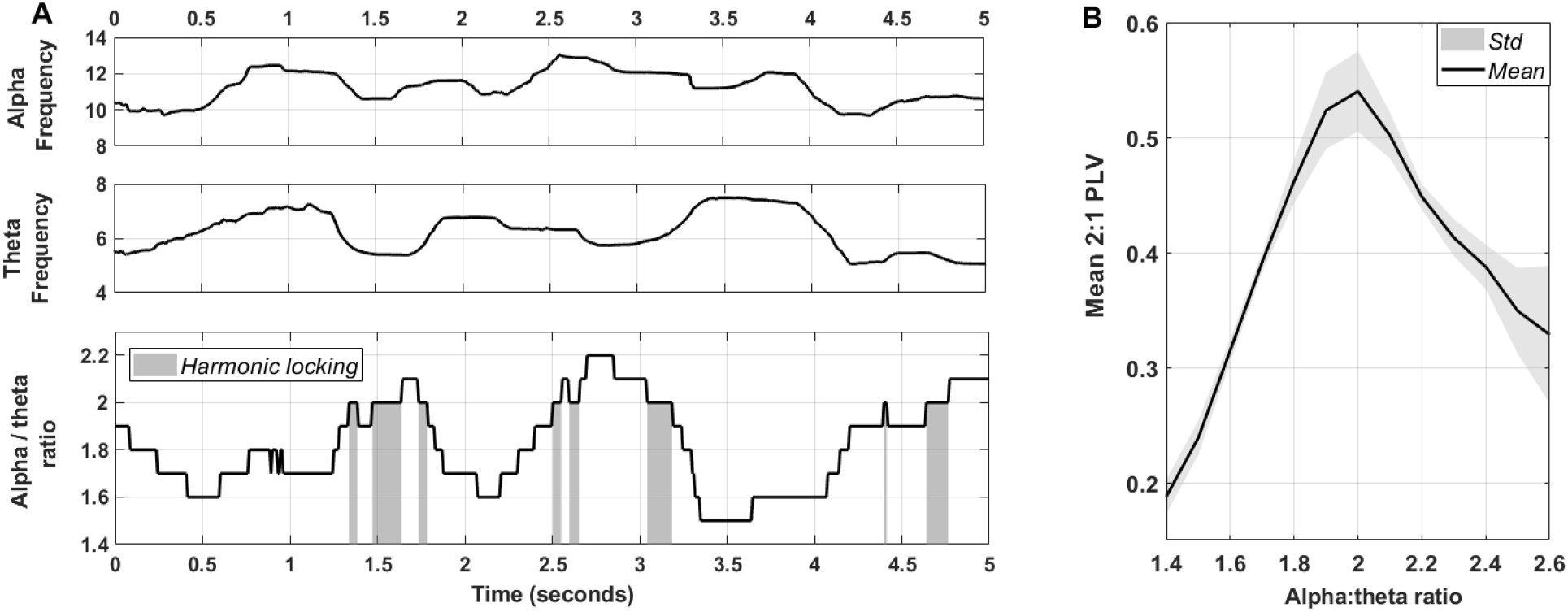
Harmonic locking estimation and its relation to alpha:theta phase synchrony. **Panel A** visualizes alpha and theta instantaneous frequencies as well as their numerical ratio over time (i.e. 5 seconds) for an exemplary subject and electrode. Grey areas indicate time points in which alpha and theta frequencies formed a 2:1 transient ‘harmonic’ relationship (i.e. harmonic locking). **Panel B** visualizes the mean 2:1 phase locking value (PLV) for distinct alpha/theta numerical ratios (recorded during rest condition) and illustrates that 2:1 phase synchrony was maximal when alpha and theta instantaneous frequencies formed a harmonic (2:1) relationship. Grey areas show standard deviation (variability between subjects).

### Cross-frequency phase synchronization estimation and its relation to harmonic locking

Since a harmonic relationship between alpha-theta rhythms mathematically enables stable and frequent phase meetings, it is expected that a greater incidence of harmonic locking corresponds to increased alpha-theta cross-frequency (2:1) phase synchronization. To illustrate this mathematical prediction in real data, alpha-theta 2:1 phase synchrony was assessed within each electrode using a phase locking metric (i.e. phase locking value), which quantifies the consistency of the phase difference between two time series. Particularly, the phase locking value is estimated as the length of a unitary vector whose angle is the instantaneous phase difference (thereby varying between 0 and 1). The phase locking value (PLV) time series was computed with a sliding window of 500 ms (1 bin step) (MATLAB code was adapted from Scheffer-Teixeira and Tort^44^).

In order to illustrate the relationship between harmonic locking and cross-frequency 2:1 phase synchrony, the average PLV was estimated for different alpha:theta ratios in 500-ms epochs per subject and across electrodes (rest condition only). Hence, Figure 1B illustrates that 2:1 phase synchrony is maximal when alpha and theta instantaneous frequencies form a harmonic (2:1) relationship.

### Statistical analysis

Condition-related differences in 2:1 alpha:theta harmonic locking were examined using the a cluster-based permutation statistical method^45^ as implemented in the MATLAB toolbox FieldTrip^46^. This statistical method controls for the type I error rate arising from multiple comparisons across electrodes through a non-parametric Montecarlo randomization. In short, data is shuffled (1000 permutation) to estimate a “null” distribution of effect size based on cluster-level statistics (sum of t-values with the same sign across spatially adjacent electrodes). Then, the cluster-corrected p-value is defined as the proportion of random partitions in the null distribution whose test statistics exceeded the one obtained for each significant cluster (p<0.05) in the original (non-shuffled) data.

In order to assess the hypothesized linear increase in harmonic locking (i.e. meditation < rest < arithmetic) the adopted test statistic (t-value) was obtained based on linear regression analysis (for dependent samples) (see ft_statfun_depsamplesregrt in FieldTrip). In addition, paired-samples t-tests were used to directly compare the incidence of harmonic locking between the conditions arithmetic–rest and rest–meditation (ft_statfun_depsamplesT in Fieldtrip). Further, independent samples F-tests (ft_statfun_indepsamplesF in Fieldtrip) were used to assess whether the difference in harmonic locking between meditation and rest varied significantly across meditation traditions.

### Secondary analysis

#### Exploring the cross-frequency ratio specificity of the condition effect

In addition to the percentage of time points displaying the aforementioned ‘harmonic’ alpha:theta cross-frequency relationship (i.e., ratio = 2:1), we computed the percentage of time points displaying other possible cross-frequency relationships to explore whether the hypothesized linear effect (i.e. meditation< rest< arithmetic) was specific to a harmonic frequency arrangement. In short, the proportion (%) of time points displaying cross-frequency ratios within a range of 1.0 to 3.5 (with a step-size of 0.1) were computed separately for each subject, condition and electrode. Then, a linear fit across the three conditions (meditation, rest and arithmetic) was estimated (ft_statfun_depsamplesregrt), separately for each electrode and cross-frequency ratio. In this case, cluster-based permutation statistics were adopted to control for multiple comparisons across electrodes and cross-frequency ratios, i.e., identifying significant clusters based on spatial (electrodes) and cross-frequency ratio adjacency.

#### Exploring instantaneous frequency modulations in the alpha and theta bands

In these analyses, we explored whether condition-related differences are also evident when modulations in instantaneous frequencies of the alpha and theta band are assessed separately. The instantaneous alpha and theta frequencies were computed per time point and rounded to the nearest integer. Then, the proportion (%) of each instantaneous frequency was calculated (i.e. 4, 5, 6, 7 and 8 Hz in the theta band and 8, 9, 10, 11, 12, 13 and 14 Hz in the alpha band). A linear fit across the three conditions (meditation< rest< arithmetic) was estimated (ft_statfun_depsamplesregrt), separately for each electrode and frequency; and cluster-based permutation statistics were adopted to control for multiple comparisons across electrodes and frequencies, i.e., identifying significant clusters based on spatial (electrodes) and instantaneous frequency adjacency. In addition, the relationship between inter-individual differences in mean alpha / theta peak frequency (i.e. instantaneous frequency averaged over time) and the incidence of harmonic locking was assessed through Spearman’s rho correlation coefficient (computed across electrodes).

#### Assessing the possibility of artefactual harmonics due to non-sinusoidal signals

Non-sinusoidal signals can produce harmonic peaks in the frequency spectra that can be wrongly interpreted as two (interacting) oscillators^47–49^. If the here reported ‘harmonic locking’ (i.e. transient 2:1 peak frequency relationship) between alpha and theta rhythms would be reflecting non-sinusoidal properties of a single rhythm, the power time series of these two harmonic frequencies are expected to be highly correlated^31,50^. To rule out this possibility, Spearman’s rank-order correlation coefficients were estimated between the power of alpha and theta frequencies that led to harmonic locking. For this purpose, the power time series in the alpha and theta frequency bands that led to harmonic locking (mean frequency +-std) were extracted per subject, electrode and condition through Short-term Fast Fourier Transform (sliding Hanning window of 1 s with 90 % overlap).

#### Identification of alpha and theta peak frequencies using the findpeaks algorithm

Previous work assessing 2:1 harmonic alpha-theta locking used the ‘findpeaks’ function implemented in MATLAB r2017b to detect transient peak frequencies within the theta (4 – 8 Hz) and alpha (8 - 14 Hz) bands^34^. To assess the consistency of the results, secondary analyses were conducted to test whether condition-related differences in harmonic locking were also evident when alpha and theta peak frequencies are estimated using this ‘findpeaks’ approach^34^. In short, transient peak frequencies in the theta (4 – 8 Hz) and alpha (8 - 14 Hz) bands were detected for each (overlapping) 1 second epochs of time-frequency transformed data (Short-term Fast Fourier Transform; 90% overlap) using the find local maxima function implemented in MATLAB r2017b. This algorithm identifies as local peaks those data samples that are larger than their two neighboring samples. When two or more peaks were detected, the peak with the highest amplitude was selected. Based on the identified alpha and theta peak frequencies, the proportion of epochs that presented an alpha:theta ratio = 2.0 (i.e. harmonic locking) was calculated per subject, electrode and condition. Condition-related differences were assessed using similar statistical analyses as described above.

## RESULTS

### Condition-related differences in 2:1 alpha:theta harmonic locking

Linear regression analysis revealed a significant positive cluster (t_cluster_ (42) = 69.68; p < 0.001) involving the majority of electrodes thereby indicating that the occurrence of harmonic locking (% of time points displaying a 2:1 relationship between alpha and theta oscillations) increased linearly from the meditation to the arithmetic condition (i.e. meditation< rest< arithmetic; Figure 2A). In addition, paired-samples t-tests revealed that the increase in harmonic locking in ‘arithmetic’ compared to ‘rest’ was more pronounced in frontal electrodes (t_cluster_ (42)= 5.65; p = 0.02) (Figure 2B) whilst the difference between ‘rest’ and ‘meditation’ was more pronounced in central (t_cluster_ (42) = 10.36; p = 0.008) and left parieto-occipital electrodes (t_cluster_ (42) = 9.02; p = 0.008) (Figure 2C).

**Figure 2.**
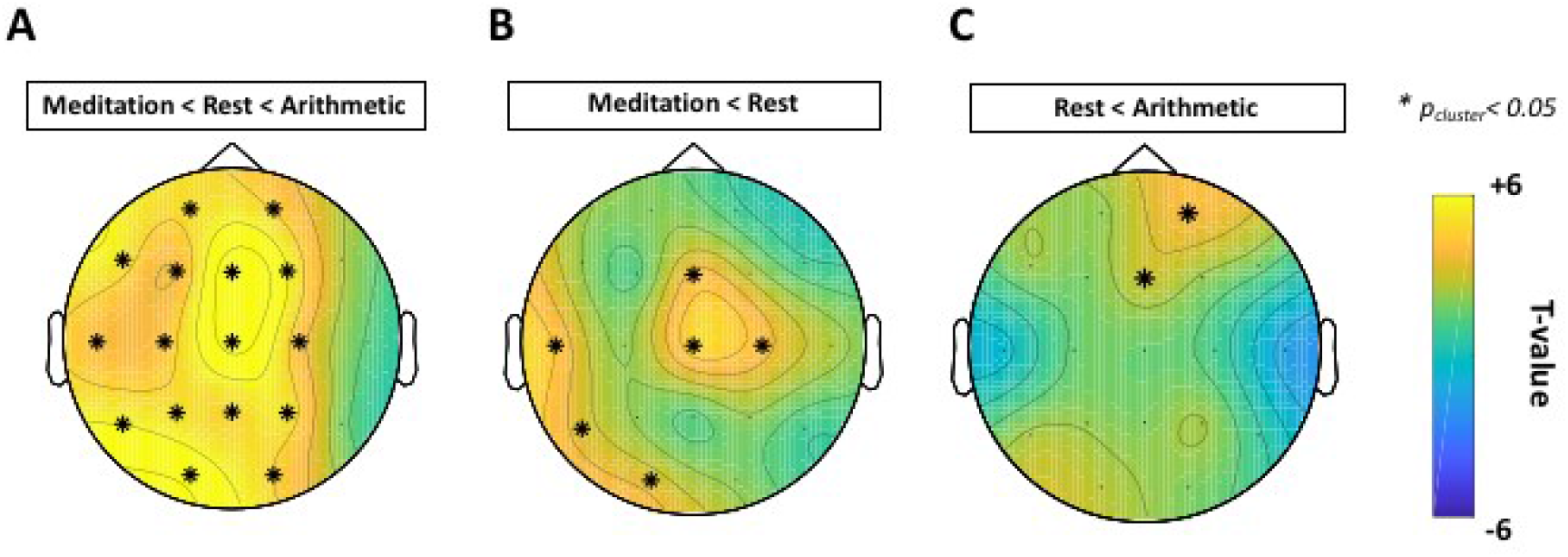
Condition-related differences in alpha:theta harmonic locking (% of time points displaying a 2:1 relationship between alpha and theta oscillations). Topographical heat maps represent t-values (panel A: linear regression; panels B-C: paired samples t-test) at each electrode and asterisks mark significant electrodes (p<0.05, cluster-based permutation analysis). As visualized in Panel A, the occurrence of harmonic locking increased linearly from the meditation to the arithmetic condition (i.e. meditation < rest < arithmetic) in a widespread cluster encompassing the majority of the electrodes. Panel B visualizes the increase in harmonic locking in rest with respect to the meditation condition (two positive clusters encompassing electrodes Fz, T3, Cz, C4, T5 and O1). Panel C visualizes the increase in harmonic locking in arithmetic with respect to the rest condition (positive cluster encompassing Fp2 and Fz electrodes).

In addition, an independent samples F-test was conducted to assess whether the difference in harmonic locking between ‘meditation’ and ‘rest’ varied significantly across meditation traditions. No significant clusters were identified, thereby indicating that the decrease in harmonic locking during meditation compared to rest did not differ significantly between meditation traditions.

### Secondary analysis

#### Exploring the cross-frequency ratio specificity of the condition-related effect

In the previous section, we assessed condition-related differences in 2:1 alpha-theta ‘harmonic locking’. In this section, we explored whether the incidence of any of the other possible cross-frequency relationships between alpha and theta frequencies also showed a linear condition-related effect (i.e. meditation – rest – arithmetic) (see *Statistical analysis*). Cluster-based permutation analysis revealed a significant positive cluster with a widespread spatial distribution covering cross-frequency ratios between 1.7 and 3.1 (t_cluster_ (42)= 643.85; p< 0.001) indicating that within this range of ratios, a linear effect was evident (i.e. meditation< rest< arithmetic), although note that the effect was maximal around the 2:1 harmonic ratio (see black box in Figure 3). In addition, this analysis also revealed a significant negative cluster (t_cluster_ (42)= −330.85; p< 0.001) with a widespread spatial distribution covering ‘lower-end’ cross-frequency ratios ranging between 1.0 – 1.5, thereby indicating a significant linear effect in the opposite direction (i.e. meditation> rest> arithmetic) for the incidence of these cross-frequency relationships (Figure 3.

**Figure 3.**
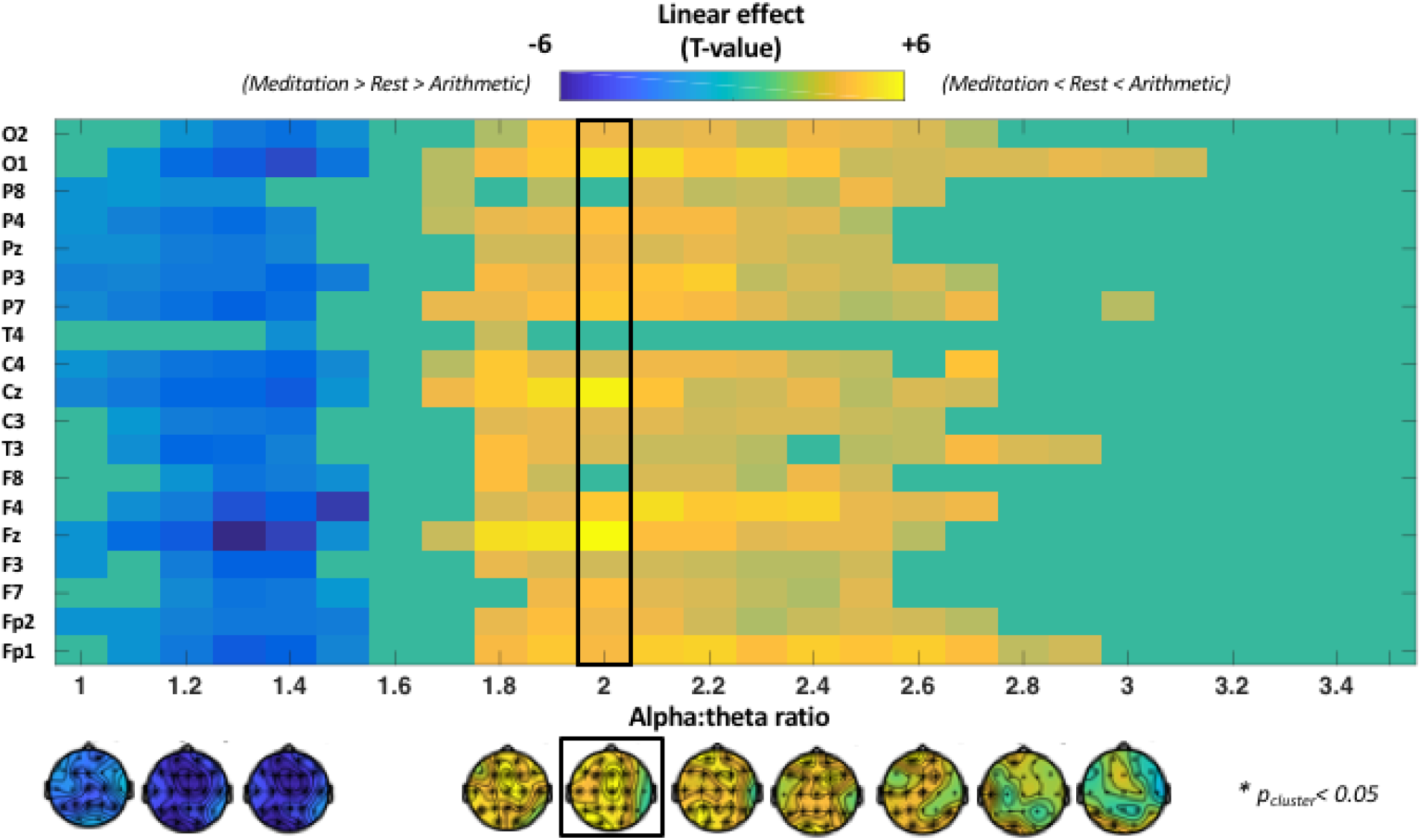
Condition-related differences in the incidence of different alpha:theta cross-frequency relationships (ratios ranging between 1.0 to 3.5). The plot visualizes the estimated linear effect (comparing meditation vs rest vs arithmetic) separately for each electrode (y-axis) and cross-frequency ratio (x-axis). A positive cluster with a widespread spatial distribution was identified for cross-frequency ratios ranging between 1.7 and 3.1, indicating that the incidence (% time points) of this range of ratios was higher in the arithmetic, compared to the rest and meditation condition (i.e. meditation< rest< arithmetic; in warm colours). In addition, negative cluster with a widespread spatial distribution was identified for cross-frequency ratios ranging between 1.0 – 1.5, indicating that for this range of ‘lower-end’ ratios, the incidence was higher during meditation, compared to rest and arithmetic (i.e. meditation> rest> arithmetic; in cool colours).

#### Exploring instantaneous frequency modulations in the alpha and theta bands

In the previous section, we explored condition-related differences in cross-frequency relationships (ratios) between instantaneous frequencies of the alpha and theta band. Here, we explored whether condition-related differences are also evident when modulations in instantaneous frequencies of the alpha and theta band are explored separately. Cluster-based permutation analysis revealed a linear increase from meditation to arithmetic (meditation< rest< arithmetic) in the incidence of instantaneous frequencies at the lower-end of the theta band, i.e., between 4 and 6 Hz (t_cluster_ = 166.81; p< 0.001); and a linear decrease (meditation> rest> arithmetic) in the incidence of instantaneous frequencies at the upper-end of the theta band, i.e., between 7 and 8 Hz (t_cluster_ = −134.49; p < 0.001). An opposite pattern was evident for alpha band frequencies, indicating a linear increase from meditation to arithmetic (meditation< rest< arithmetic) in the incidence of upper-end alpha frequencies, i.e., between 11 and 14 Hz (t_cluster_ = 116.43; p < 0.001); and a linear decrease (meditation> rest> arithmetic) in the incidence of lower-end alpha frequencies, i.e., between 8 and 10 Hz (t_cluster_ = −103.16; p < 0.001) (Figure 4, right panel). Together, this pattern of results indicates a *deceleration* of instantaneous theta frequencies in the arithmetic, compared to the rest and meditation conditions (i.e., higher/lower incidence of lower/upper-end theta frequencies); and an *acceleration* of instantaneous alpha frequencies (i.e., higher/lower incidence of upper/lower-end alpha frequencies) (Figure 4, left panel).

**Figure 4.**
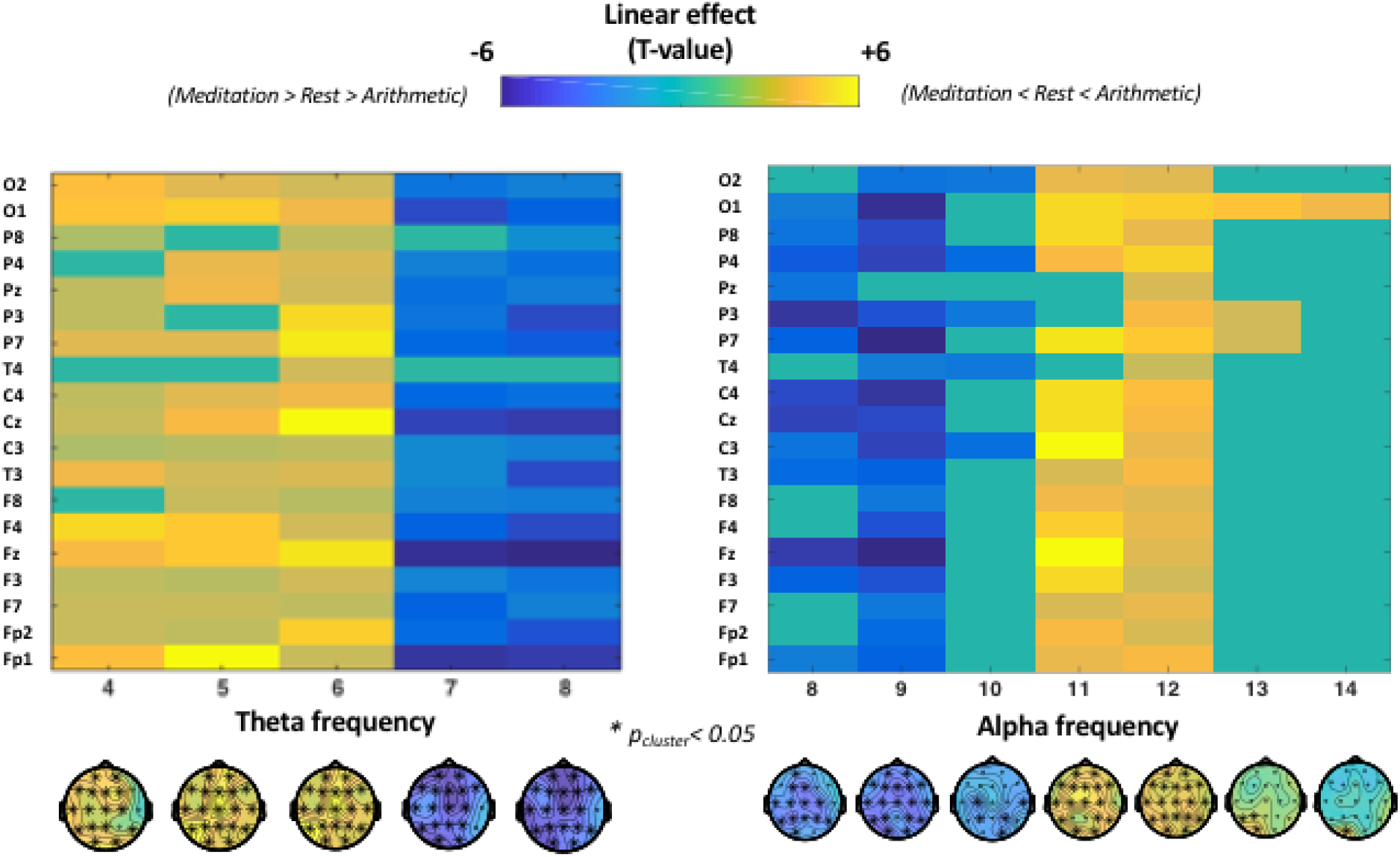
Condition-related differences in the instantaneous frequencies of the alpha and theta bands. The plots visualize the estimated linear effect (comparing meditation vs rest vs arithmetic) separately for each electrode (y-axis) and frequency (x-axis). As visualized in the left panel, the frequency of theta oscillations decelerated in arithmetic task (compared to rest and meditation conditions) with a linear increase in the occurrence of instantaneous frequencies between 4 and 6 Hz (positive cluster in warm colours; i.e. meditation< rest< arithmetic) and a linear decrease in the occurrence of instantaneous frequencies between 7 and 8 Hz (negative cluster in cool colours; i.e. meditation> rest> arithmetic). On the other hand, alpha oscillations (right panel) were shown to accelerate during arithmetic task (compared to rest and meditation conditions) with a linear increase in occurrence of instantaneous frequencies between 11 and 14 Hz (positive cluster in warm colours; i.e. meditation< rest< arithmetic) and linear decrease in the occurrence of instantaneous frequencies between 8 and 10 Hz (negative cluster in cool colours; i.e. meditation> rest> arithmetic).

Furthermore, inter-individual differences in the increased incidence of harmonic locking in the arithmetic compared to the meditation condition (i.e. harmonic-locking(arithmetic) - harmonic-locking(meditation) difference score) was proportional to alpha frequency acceleration (i.e. mean-alpha-frequency(arithmetic) - mean-alpha-frequency(meditation) difference score) (rho= 0.82; p<0.01) and theta frequency deceleration (i.e. mean-theta-frequency(arithmetic) - mean-theta-frequency(meditation)) (rho= −0.79; p< 0.01).

#### Assessing the possibility of artefactual harmonics

Spearman’s rank-order correlations between the time courses of alpha and theta power at the frequencies that led to harmonic locking were generally small and not significant (mean r-value (across subjects and electrodes) = 0.07; p= 0.12). As such, it is anticipated that the identified harmonic relationships between alpha-theta frequencies are unlikely to reflect harmonics induced by non-sinusoidal properties of a single oscillator.

#### Condition-related differences when using the findpeaks algorithm

Secondary analyses were performed to assess condition-related differences in harmonic locking when transient alpha and theta peak frequencies were estimated using the findpeaks algorithm^34^. Overall, a qualitatively similar pattern of results was revealed, indicating that harmonic locking (% of time points displaying a 2:1 relationship between alpha and theta oscillations) increased linearly from the ‘meditation’ to the ‘arithmetic’ condition (i.e. meditation< rest< arithmetic; t_cluster_ (42)= 7.24; p= 0.004; electrodes P3, P4 and Pz). Direct comparisons between conditions also revealed a trend-level increase in harmonic locking in the ‘arithmetic’ compared to the ‘rest’ condition (t_cluster_ (42)= 2.74; p= 0.07; electrode Fp2), and in the ‘rest’ compared to the ‘meditation’ condition (t_cluster_(42)= 2.99; p= 0.057; electrode O2).

## DISCUSSION

This study assessed whether the incidence of an EEG cross-frequency arrangement that facilitates cross-frequency phase synchronization between alpha and theta rhythms increases linearly from a meditative to an active cognitive processing state. In line with our hypothesis, we showed that *harmonic locking* (% of time in which the instantaneous frequencies of alpha and theta rhythms arrange in a 2:1 harmonic position) increased linearly from meditation to arithmetic task (meditation < rest < arithmetic) in a sample of 43 highly experienced meditators. Although the linear effect had a widespread spatial distribution, the increase in harmonic locking during rest compared to meditation was more pronounced in central and left temporoparietal electrodes while the increase in arithmetic compared to rest displayed a frontal distribution.

Since neural oscillations at different frequencies have been shown to play distinct functions^51–53^, it can be expected that this information has to be somehow integrated in the brain. Recent research suggests that the integration of distinct neural oscillations is implemented via cross-frequency phase synchronization as this is the only form of cross-frequency coupling that allows rigorous spike-time relationships between neural assemblies^31,50,54,55^. Crucially, strict phase synchronization is only possible for oscillators that form a harmonic relationship^30^. Based on this mathematical reality, it has been recently posited that transient changes in peak frequencies within different brain rhythms^24,25,43,56^ could reflect a key mechanism for coordinating cross-frequency coupling and decoupling in the brain (for extent see Klimesch, 2012, 2013, 2018). In this view, when information integration is needed to accomplish a specific task, the peak frequency arrangement of task-relevant rhythms is anticipated to change into a harmonic relationship in order to facilitate the communication between their underlying neural networks via cross-frequency phase synchronization. In line with this notion, it has been recently shown that the incidence of transient harmonic relationships between alpha and theta rhythms is tightly linked to cross-frequency phase synchronization, increases significantly during an arithmetic task and predicts behavioural performance^34^. In the current study, we extend these findings by showing that, in experienced meditators, harmonic locking between alpha and theta rhythms not only increases during active cognitive processing (arithmetic task condition) but also significantly decreases during meditation compared to rest, thereby providing initial support to the notion that meditative states are characterized by a reduction in cross-frequency interactions between alpha and theta rhythms.

The interaction between alpha and theta rhythms has been shown to be crucial in cognitive tasks that involve memory and executive control. Several studies suggest that the alpha rhythm encompasses the memory component (storage and retrieval of information) whilst theta is more important for the manipulation and coordination of stored information (i.e. executive component)^13–16^. Similar to laboratory cognitive tasks, self-generated thought during rest is also expected to involve some memory and executive functions (i.e. retrieval and manipulation of *internal* information). Since experienced meditators have reported not only reduced frequency of mind wandering but also attenuated semantic association to their contents during meditation practice^5–9^, we hypothesized that retrieval, retention, and manipulation of information (and therefore, the interplay between alpha and theta rhythms) would be reduced during meditation compared to rest. In line with this hypothesis, we show that the incidence of a frequency arrangement that enables alpha:theta interactions via cross-frequency phase synchrony increases linearly from meditation to arithmetic task (meditation< rest< arithmetic). Hence, our results suggest that self-generated thought during rest and controlled processes during cognitive tasks could share some neurocognitive mechanisms that are encompassed in alpha-theta cross-frequency dynamics.

Although harmonic locking between alpha and theta rhythms was shown to increase linearly from meditation to an arithmetic task (i.e. meditation< rest < arithmetic) in the majority of the electrodes (see Figure 2A), the comparison meditation vs rest and rest vs arithmetic revealed two specific topographical distributions (Figure 2B-C). Previous studies have suggested that self-generated thought and controlled processes (such as active arithmetic task performance) can recruit different networks, or the same networks but to a different extent. Particularly, while controlled cognitive processes have been consistently linked to a ‘superordinate cognitive control network’ involving dorsolateral prefrontal, anterior cingulate, and parietal cortices^58^, self-generated thought has been shown to involve (mainly) default mode network activity (DMN)^59^. Accordingly, DMN activity and connectivity have also been shown to decrease during meditation in experienced meditators^60,61^. Thus, in the light of previous literature, it seems plausible that the here obtained topographical distribution for the comparisons between arithmetic – rest and rest – meditation could represent executive control network and DMN respectively (or at least, different subdivisions within these two networks^62,63^. Nevertheless, spatial localization of harmonic locking between alpha and theta rhythms was beyond the scope of our study and future studies with better spatial resolution (e.g. high-density EEG, combined EEG-fMRI) are warranted to firmly disentangle the sources of these apparent spatial differences.

In addition to our main analysis, we performed two exploratory analyses in which we assessed whether the reported linear effect in harmonic locking i) extended to the incidence of other cross-frequency ratios and ii) was accompanied by instantaneous frequency modulations within alpha and / or theta bands separately. These analyses revealed that during an arithmetic task (compared to rest and meditation) alpha and theta rhythms separated from each other (acceleration of alpha/ deceleration of theta) thereby increasing the incidence of cross-frequency ratios approximating the harmonic (i.e. greater occurrence of transient alpha peaks around 11-12 Hz and theta peaks around 4-6 Hz). On the other hand, during meditation (compared to rest and arithmetic task) alpha and theta instantaneous frequencies tended to converge in the upper theta / lower alpha band (i.e. 7 – 9 Hz), thus increasing the incidence of cross-frequency ratios between 1.0 and 1.5. These observations are in line with previous evidence showing that effortful cognition is accompanied by an acceleration alpha peak frequency^24^, deceleration of theta peak frequency^34,64^ and increased alpha:theta cross-frequency coupling^20,21,23,34^ whilst meditative states are accompanied by power increases in the lower alpha / upper theta band Based on our results and previous literature, we speculate that when cognitive demands increase, alpha and theta rhythms separate from each other in the frequency spectrum (alpha accelerates, theta decelerates) to exert their respective functions (i.e. upper alpha = retrieval and storage information; lower theta = manipulation of information^18,19,22^, while, at the same time, allowing their interaction by the formation of transient harmonic relationships (i.e. enabling cross-frequency phase synchronization for short periods)^30,32^. On the contrary, we speculate that when information is not to be retrieved, stored and manipulated (like in meditative states that aim at achieving ‘mental emptiness’), alpha and theta rhythms tend to converge in upper theta / lower alpha band thereby avoiding the exertion of their respective functions (by oscillating at a different frequency) and their interaction (via a non-harmonic cross-frequency arrangement).

Together, our results suggest that the neural correlates of meditative states can be characterized in opposition to active cognitive processing when studied from an EEG cross-frequency dynamics perspective. Specifically, the incidence of transient harmonic relationships between alpha and theta oscillations (enabling synchrony and therefore communication between the neural networks underlying these rhythms) was shown to increase linearly from meditation to an arithmetic task (meditation< rest< arithmetic) in a sample of 43 highly experienced meditators. Our interpretation of these results is that some neurocognitive mechanisms that are present during both self-generated thought and controlled cognitive processes (i.e. the integration between the memory and executive components of cognition via alpha:theta cross-frequency coupling) are minimized during meditative practices aimed at achieving ‘mental emptiness’. From a translational perspective, these insights may hold potential for the development of novel neurofeedback protocols aimed at facilitating the achievement of meditative states by modulating alpha-theta cross-frequency dynamics.

## AUTHOR CONTRIBUTIONS

S.T. collected the data. P.F. pre-processed the data. J.R. formulated the hypothesis, performed the analysis and drafted the first version of the manuscript. J.R. and K.A. wrote the manuscript. All the authors participated in the review and approved the final manuscript.

## ADDITIONAL INFORMATION

### Competing Interests

The authors declare no competing interests.

### Data Availability

The data set analysed in the current study is available upon request.

